# Drug Proarrhythmic Evaluation in a High Throughput Cardiac New Approach Methodology

**DOI:** 10.64898/2026.05.11.722965

**Authors:** Verena Charwat, Adrian Ramirez, Karoline H. Jæger, Brennan Kandalaft, Henrik Finsberg, Brian Siemons, Aslak Tveito, Kevin E. Healy, Samuel T. Wall

**Affiliations:** Organos Inc, Berkeley, CA, USA; Johannes Kepler University, Department of Pathophysiology, Linz, Austria; Simula Research Laboratory, Oslo, Norway; Departments of Bioengineering, and Materials Science & Engineering, University of California Berkeley, Berkeley, CA, USA

## Abstract

**Background and Purpose:** Cardiotoxicity is a major cause for drug failure throughout the drug development process, with particular concern for action potential prolongation and arrhythmia. Hence, such liabilities are heavily considered during the early phases of drug design to prevent dangerous compounds from progressing. New approach methodologies (**NAMs**) that efficiently examine this risk early in the discovery pipeline should help streamline drug development programs. We developed a cardiac NAM, a 384-well open bath platform consisting of cardiac tissue derived from human induced pluripotent stem cell (**hiPSC**)-derived cardiomyocytes, enabling high-throughput drug screening while maintaining the structural and functional complexity of 3D cardiac micromuscles.

**Methods:** We dramatically increased throughput without compromising physiological relevance provided by the 3D micromuscle structure. Our 384-well open bath high-throughput platform allowed evaluation of multiple compounds at a time, enabling us to study the CiPA (comprehensive in vitro proarrhythmia assay) drug panel for proarrhythmia screening. We obtained phenotypic fingerprints of all 28 compounds (9 low, 11 intermediate, and 8 high arrhythmia risk; https://cipaproject.org) in dose-escalation studies around their respective clinical concentrations. The analysis was augmented with an *in silico* pipeline that used phenotypic biomarkers to invert data into a mathematical model of cellular currents to infer which ion channels were affected upon drug exposure, and then trained a ML model to predict channel block.

**Results and Conclusions:** We found accurate detection of arrhythmic potential for most of the compounds, and the *in silico* model inversions were consistent with published values of compound channel block. All the high risk compounds showed action potential duration (**APD**) prolongation coupled with either action potential abnormalities, early afterdepolarizations (**EADs**), or beat cessation. For the intermediate risk group, 9 out of 11 compounds caused APD prolongation alone or in combination with EADs while 2 others showed either beat cessation or beat rate change. Augmentation of APD analysis with detailed biophysical modeling and ML tools provided meaningful insight into the mechanisms involved in APD changes. Overall, our cardiac NAM allowed for fast and relevant screening for mechanistic understanding of APD prolongation and proarrhythmic activity, at massively increased throughput compared to other 3D micromuscle models.

**Summary:** Cardiotoxicity testing is critical in drug development to prevent arrhythmogenic side effects. Current stringent regulations have greatly reduced market withdrawals; however, these strict evaluations often lead to costly late-stage failures and loss of promising candidates as false positives. We developed a cardiac new approach methodology (NAM), a 384-well open bath cardiac micromuscle platform created from hiPSC-derived cardiomyocytes, enabling high-throughput drug screening while maintaining the structural and functional complexity of 3D cardiac micromuscles. Using the comprehensive in vitro proarrhythmia assay (CiPA) drug panel, we validated the system to accurately detect proarrhythmic potential. Our assay provided phenotypic fingerprints based on mechanical and electrophysiological biomarkers. Integration with computational modeling offered insights into multi-ion channel effects (MICE). Particularly, we identified sodium channel block contributions as a significant factor for poor risk prediction based on traditional parameters. The combined experimental and computational platform can enhance early drug screening, thereby reducing late-stage failures and promoting the progression of low-risk compounds with complex electrophysiological profiles.

## Introduction

Minimizing and eliminating off-target toxicity is still a primary focus of drug discovery pipelines, where significant resources are invested to ensure that clinical candidates are free of hidden, serious adverse effects. Cardiovascular toxicity in particular is one of the largest concerns in novel drug safety testing, in large part due to a history of compounds that have made it to market, only to be removed at great cost due to deadly adverse cardiac events, such as arrhythmia, Torsades de Point (TdP).^1^ Due to this history, a large effort has been made to ensure that drugs that progress through development pipelines lack arrhythmogenic cardiac side effects. This effort has largely paid off, with no market removals of drugs over the past decade due to TdP.

However, this success has come at a substantial cost to the drug development pipeline. It requires extensive *in vitro* and *in vivo* testing, combined with preclinical and early clinical evaluation, to fully establish the safety profiles of compounds. These tests generally include studies on cardiomyocyte membrane ion channel interaction, especially the human ether-a-go-go related gene protein (hERG), heavily implicated in arrhythmia risk, as well as, where warranted, further studies on the QT interval in canines and later monitoring in humans. While this pipeline has been generally successful for use on advanced candidates, the overall workflow is cumbersome, expensive, and most importantly, possibly too restrictive. In particular, the historical focus on excluding hERG interacting molecules, which while successful at eliminating drug induced QT prolongation, is clearly limiting, as QT prolongation is not uniquely predictive of cardiac risk,^2, 3^ and safe compounds with hERG block are used clinically.^4^

Moreover, compounds with multi-ion channel effects (**MICE**) are particularly challenging to classify regarding their arrhythmogenic potential. Most prominently, calcium channel blockage (partially) counteracts hERG-block effects on the cellular action potential.^5^ Moderate differences in the ion channel expression profile in preclinical models compared to the human *in vivo* situation can skew safety test results and lead to false negative or false positive predictions. As the exclusion of potentially useful safe drugs hurts both drug developers and patients, this limitation has stimulated the development of better tools, such as NAMs, and guidelines that can be useful for more reliable classification of cardiovascular risk in drug compounds.

Over the past decade, proarrhythmic evaluation using hiPSC-derived cardiomyocytes (**hiPSC-CM**) has emerged in a myriad of assays. Using standard cell culture techniques, these assays can be run at scale, resulting in greater throughput than the traditional safety testing in animal models. The ability of these methods to detect changes in both electrophysiology and mechanics provides a more holistic phenotypic understanding of how a compound may alter the cardiovascular system compared to other methods, such as ion channel interaction tests. In particular, the CiPA initiative, which was launched in 2013 as a dedicated effort to improve the arrhythmia risk classification using a combination of approaches including hiPSC-CM models, has found that these cells can largely be used to predict risk in known compounds, with some caveats.^5, 6^ The limitations identified with proarrhythmia risk testing in hiPSC-CM models include the cells’ relative immaturity, variability between hiPSC-CM lines, lack of standardization, complexity of drug effects (multi-channel effects, compounding factors such as comorbidities) and oversensitivity (false positive results).^7^

One potential way to improve on these efforts is to change from 2D cultures to 3D microtissues, as extensive evidence indicates that 3D cardiac tissue models are more physiologically predictive. The 3D microenvironment enhances cell-cell and cell-matrix interactions, leading to improved structural, electrophysiological, and contractile maturation. Vunjak-Novakovic *et al.*^8, 9^ demonstrated that mechanical and electrical conditioning in 3D tissues yields more adult-like cardiac phenotypes. Likewise, Huebsch *et al.*^10^ showed that 3D microtissues more accurately recapitulate pharmacologic effects, such as Verapamil’s effect on contractility, than 2D systems. Similar findings from Mannhardt *et al.*^11^ confirm the superior predictive power of 3D engineered tissues for drug response and proarrhythmic risk.

A challenge is maintaining the throughput capacity of 2D cultures with a matured 3D tissue format. Here we present a proarrhythmic risk platform that is based on an industry standard 384 well plate layout with custom substrates for 3D cardiac micromuscle formation. It allows for electrophysiological and mechanical readouts of ∼ 80 wells containing up to 320 micromuscles in under 1 hour. Phenotypic fingerprinting is achieved for each tissue by direct assessment of the experimental data (e.g. action potential duration increase), and post-processing via computational models and model trained AI algorithms. We found that this *in vitro/in silico* layered approach produced a high degree of insight into compound cardiotoxicity, with dose dependent APD changes consistent with known drug effects, with computational modelling providing information on drug-induced ion channel block and effects on the mechanic intracellular machinery.

## Results

### Cardiac NAM – a high throughput platform for cardiac micromuscle for proarrhythmia screening

We created cardiac micromuscles in a 384 well plate format using custom substrates patterned with micron scale troughs (i.e., microtroughs) (**Fig. 1A-F**). After loading hiPSC-CM single cell suspensions, the cells spontaneously formed micromuscles without the need for additional matrix material (**Suppl Fig. 1**). While cell layers growing on flat thermoplastic elastomer (TPE) surfaces showed disorganized cell orientation (**Fig. 1G**, top), the cells in the microtroughs formed a micromuscle spanning between the two adjacent sets of anchor pillars (**Fig. 1E**). Sarcomeric structures were stained and striations confirmed uniaxial cell elongation in the 3D micromuscles (**Fig. 1G**, bottom). Cross-sections reconstructed from confocal Z-stacks revealed a true muscle fiber geometry containing about two cell layers for every 10µm of micro tissue thickness (**Fig. 1H**). Spontaneous beating activity started around day 2 to 3 after loading and cells continued to condense into microtissues for about a week. After that the micromuscles remained stable with spontaneous beat rates of 0.59 *±* 0.14 Hz and cAPD80_0_ of 376 *±* 47 ms for up to two weeks.

**Figure 1:**
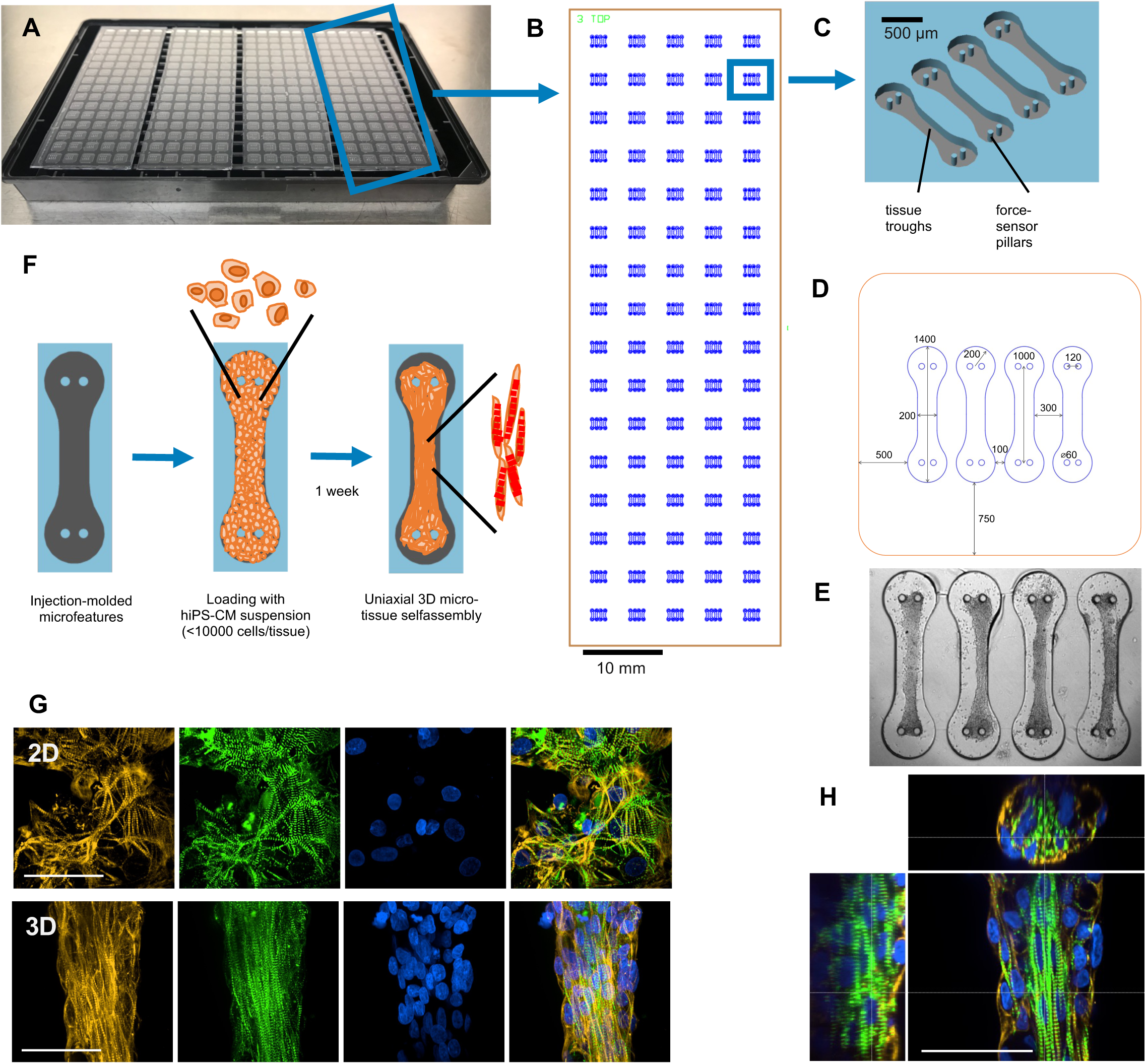
High throughput platform for cardiac micromuscle formation. **A**) Plate assembly of a 384 well plate with 4 strips of micropatterned TPE. **B**) Design of a single strip of TPE with an array of 16x5 micropatterned areas (blue) containing 4 troughs per well. **C**) 3D rendering showing the 4 micromuscle troughs in the substrate of each well **D**) Dimensions of the micromuscle troughs. The orange rim indicates the size of a square 384 well. **E**) Brightfield image of cardiac micromuscles formed in 4 troughs. **F**) Concept of cell loading and tissue formation in a micromuscle trough. **G**) Maximum intensity projection of immunofluorescence confocal Z stacks (17µm total thickness, 34 slices) of cardiomyocytes on the flat TPE substrate (top) and a cardiac micromuscle spanning between anchor pillars (pillars located outside of the imaged region) (bottom). Samples stained for sarcomeric structures (cTNNT, orange; F-actin (Z-discs), green), nuclei (DAPI, blue) and overlay. H) Reconstructed cross-sections along the x, y and z axes of a confocal immunofluorescence image stack of the cardiac micromuscle. Scalebars in G) and H) represent 50µm.

### Experimental assessment of proarrhythmic markers

Drug induced APD prolongation is a main criterion for proarrhythmic risk flagging of new compounds. All drugs were tested in a dose-escalation manner on the 3D micromuscles, and brightfield and fluorescence images and videos were recorded for phenotypic fingerprinting of APD and other biomarkers (**Fig. 2A**). Most compounds in our data set were correctly divided into risk groups based on APD prolongation in relation to their respective C_max_ (**Fig. 2B**). Low risk compounds (green) typically showed no APD increase, or even APD shortening. Datapoints remain below the cut-off for APD increase over all tested doses. Most high risk compounds (red) have datapoints in the upper left quadrant showing strong APD increase at doses below or around C_max_ (**Fig. 2B**). Intermediate risk compounds (yellow) were the most variable group, mostly showing APD increase at doses beyond 10 or 30-fold C_max_ (upper right quadrant) (**Fig. 2B**). Similar studies have been performed by multiple groups with largely matching results concluding that hiPSC-CM models are generally suitable for proarrhythmia risk prediction with come caveats and limitations.^7^

**Figure 2:**
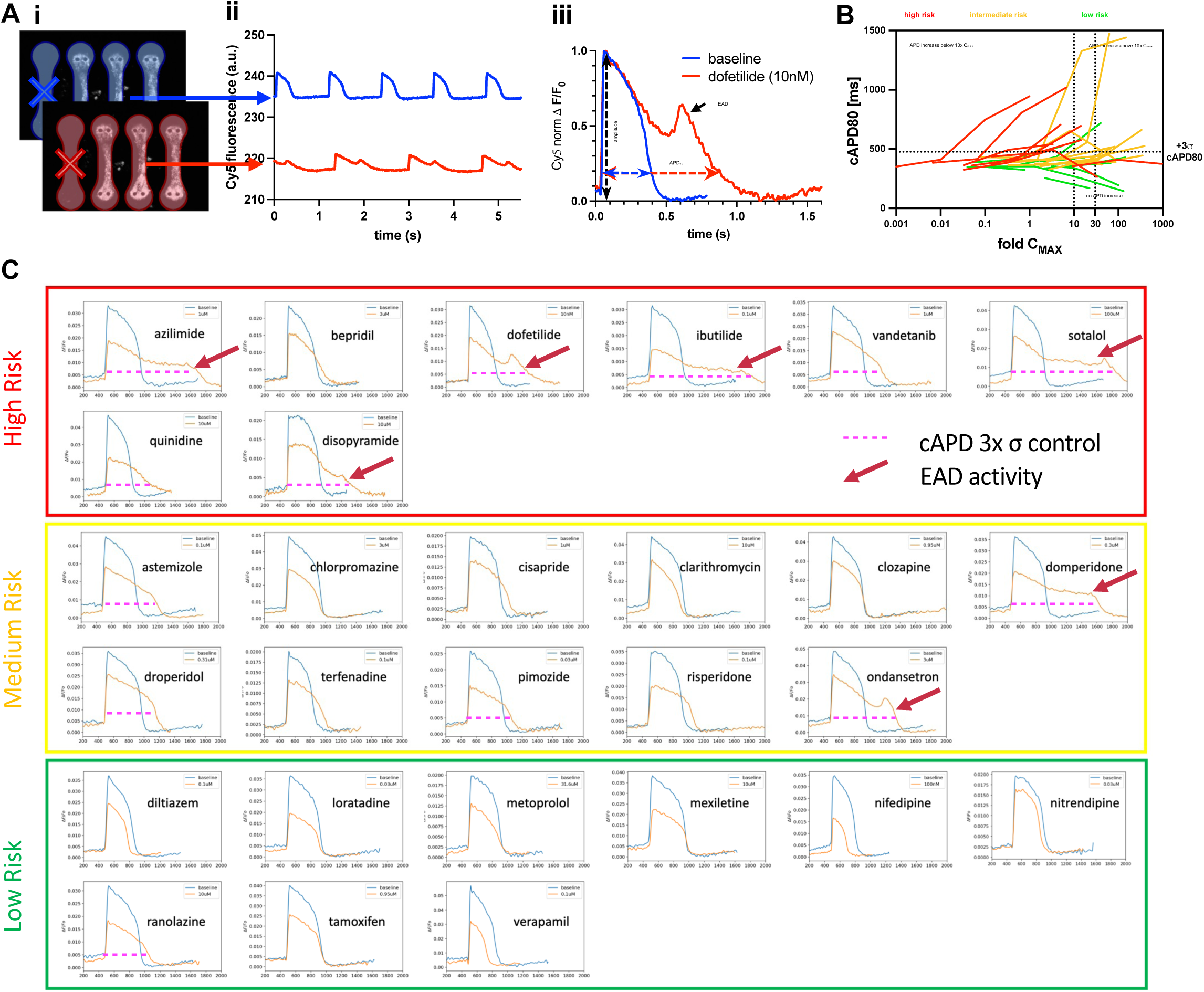
Experimental assessment of pro-arrhythmic markers. **A**) Data acquisition and processing schematic for voltage traces: Videos of PhoS-1 stained tissues are acquired in the Cy-5 channel after exposure to each drug dose. (**i**) Each video contains recordings of all 4 tissue troughs in a well. A mask is applied (colored areas) to distinguish different tissues for data collection. (**ii**) Intensity-time curves are extracted via automated scripts and low quality traces are removed from further analysis based on exclusion criteria (e.g., crossed-out empty troughs in **i**). Traces are processed to generate background subtracted averaged beat traces from which parameters such as APD at 80% peak amplitude (APD_80_) are computed. (**iii**) Beat irregularities, mainly early afterdepolarizations (EADs), were identified manually. **B**) Beat rate corrected APD values (cAPD80) are plotted against the clinical maximum plasma concentration (C_max_) for all 28 compounds. Compounds are color coded: red for high risk, yellow for intermediate risk, and green for low risk. The horizontal cut-off bar (dotted line) indicates an APD increase more than 3x the standard deviation of all baselines. The vertical lines mark safety margins (10 and 30 - fold C_max_) relevant for the drug development pipeline. cAPD80 data was only plotted if values could be extracted from at least 2 traces. Beat cessation and/or strong arrhythmias resulted in inability to accurately determine the cAPD80. **C**) Representative voltage traces from all 28 tested compounds classified by risk. Beat averaged AP waveform, measured using a voltage sensitive dye, is depicted for a control acquisition and for the highest measured concentration where beating was still observed. Absolute values are depicted, with a drop in fluorescence magnitude due to photobleaching over multiple acquisitions. Significant changes in APD (cAPD80 increase more than 3x standard deviation σ of the controls) are marked with red dotted lines. Red arrows indicated EAD occurrence.

After this coarse quality check of our model, we performed a more detailed analysis for each compound. We plotted dose-dependent cAPD80 changes together with any observation of EADs, a prominent *in vitro* arrhythmia indicator. **Fig. 2C** shows representative voltage traces for all 28 compounds at a single dose. EADs are indicated by red arrows and traces with cAPD80 increases greater than 3 times the standard deviation of the controls are marked with pink dashed lines. **Fig. 3** summarizes the action potential data across all compounds, concentrations, and replicates. For all compounds our APD analysis is consistent with results observed in the literature.^6, 7, 12-14^ As a main risk predictor (*predictor A*), we considered either the occurrence of EADs (red bars) or increased cAPD80 values (red symbols) above a threshold value. The threshold was defined as the average cAPD80 value of all baselines in the study + 3 times their standard deviation (horizontal dotted line in **Fig. 3**). Additionally, a less strict risk criterion (*predictor B*) was defined as cAPD80 values that were: a) significantly different from their own baseline values; and, b) the cAPD80 prolongation was larger than 2 times the standard deviation of the respective drug dose (yellow symbols). All except one of the high risk compounds met criteria of *predictor A* at doses around or slightly above their clinical C_max_. The only exception was bepridil, which only hit *predictor B* criteria at doses almost 94-fold C_max_. For the medium risk group, most compounds met either *predictor A* (astemizole, cisapride, clarithromycin, domperidone, droperidol, pimozide, ondansetron) or at least *predictor B* (cisapride, terfenadine, risperidone). Notably, the observed proarrhythmic risk markers were typically only met at doses much higher than C_max_ compared to the high risk group. Only 2 compounds of the medium risk group were not classified for arrhythmia risk (chloropromazine and clozapine). Most of the low risk compounds did not meet any of the risk prediction criteria. Exceptions were mexiletine, ranolazine and tamoxifen. For tamoxifen, EADs were detected in a low percentage of tissues at the highest measured dose (143-fold C_max_). Ranolazine met predictor A for APD prolongation at only 5.1-fold C_max_. Similarly, mexiletine showed EADs and beat rate increase a just 4-fold C_max_.

**Figure 3:**
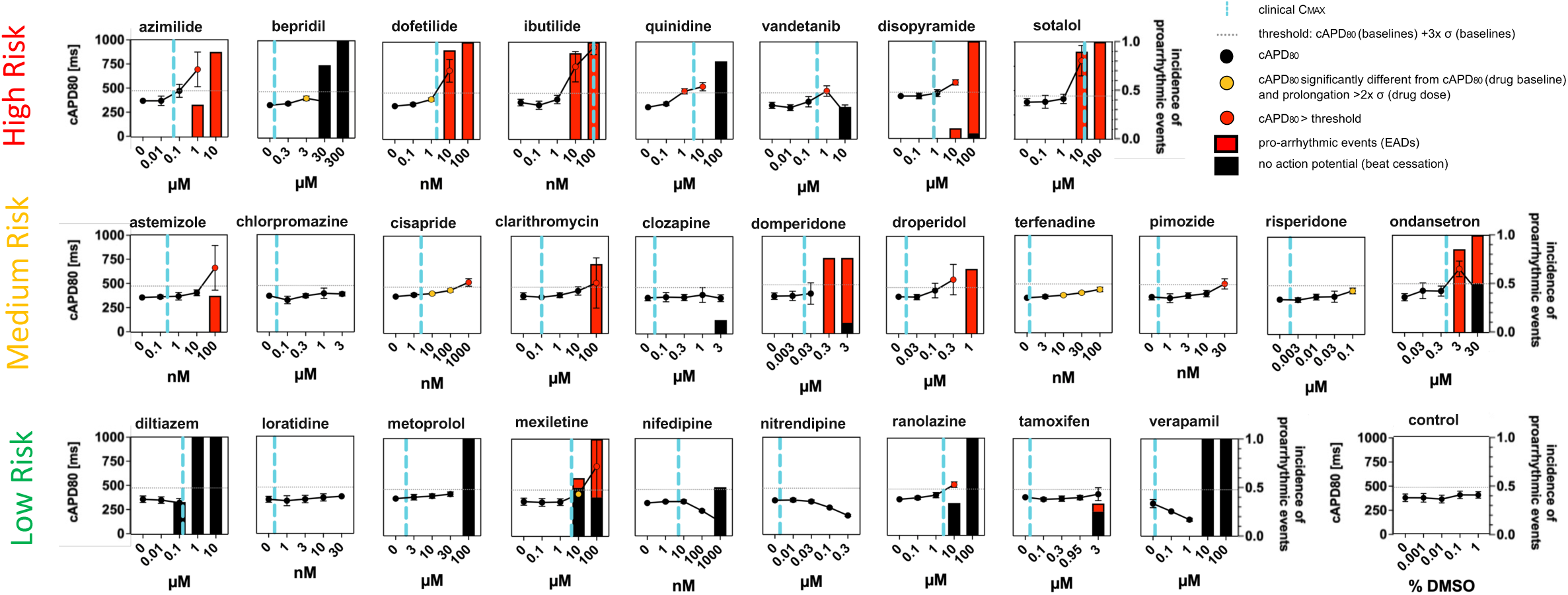
Action potential data for 28 CiPA compounds. cAPD80 values (circles + lines) and incidence of proarrhythmic events (bars) of 5-step dose response curves were plotted for all 28 CiPA compounds. The red portion of each bar shows proarrhythmic events and the black portion shows absence of a detectable AP. Error bars show standard deviation of cAPD80; σ is the standard deviation of cAPD80 values from all baseline recordings. Data from a minimum of n=5 (and up to n=21) tissues was included for each compound. Dashed vertical lines (blue) represent the compound’s clinical maximum plasma concentration (C_max_). Dotted horizontal lines represent a threshold for cAPD80 increase defined as baseline cAPD80 + 3*σ. cAPD80 values above the threshold line have red symbols. cAPD80 values that are significantly different from the compound’s baseline and have an increase of least 2x standard have orange symbols. cAPD80 values above the chart range (>1000ms) are not plotted. cAPD80 data was only plotted if values could be extracted from at least 2 traces. Beat cessation and/or strong arrhythmias resulted in inability to accurately determine the cAPD80.

### Computational model analysis of experimental data

The computation results provide additional and critical contextualization to these results. We present data fit to models of channel block (dashed purple line) to a wide range of previously published measurements of drug channel interaction (gray lines) (**Fig. 4A**). Modeling results based on experimental biomarkers are broadly consistent with literature values, demonstrating that theoretical biophysical representations are capable of accurately capturing known single and multi-ion channel effect (MICE). The only high risk compound that did not produce a *predictor A* result, Bepridil, showed both calcium and sodium blocks in the model. Together, these two blocks can mitigate action potential prolongation. For the intermediate group, again hERG block was observed for most compounds, even for compounds that did not see substantial experimental APD prolongation or EAD behavior (chloropromazine and clozapine). In the case of chloropromazine, as with bepridil, there are concurrent changes to both calcium and sodium currents, while clozapine shows a calcium block, demonstrating how the balance of these channel blocks can dramatically alter compound risk classification. For the low risk compounds, a clear pattern emerged of low hERG block near the C_max_, or a strong simultaneous calcium block. Interestingly, by analysis of action potential dynamics alone, several ‘safe’ compounds were flagged, namely mexiletine, ranolazine, and tamoxifen, compounds having the common feature of a sodium block.

**Figure 4:**
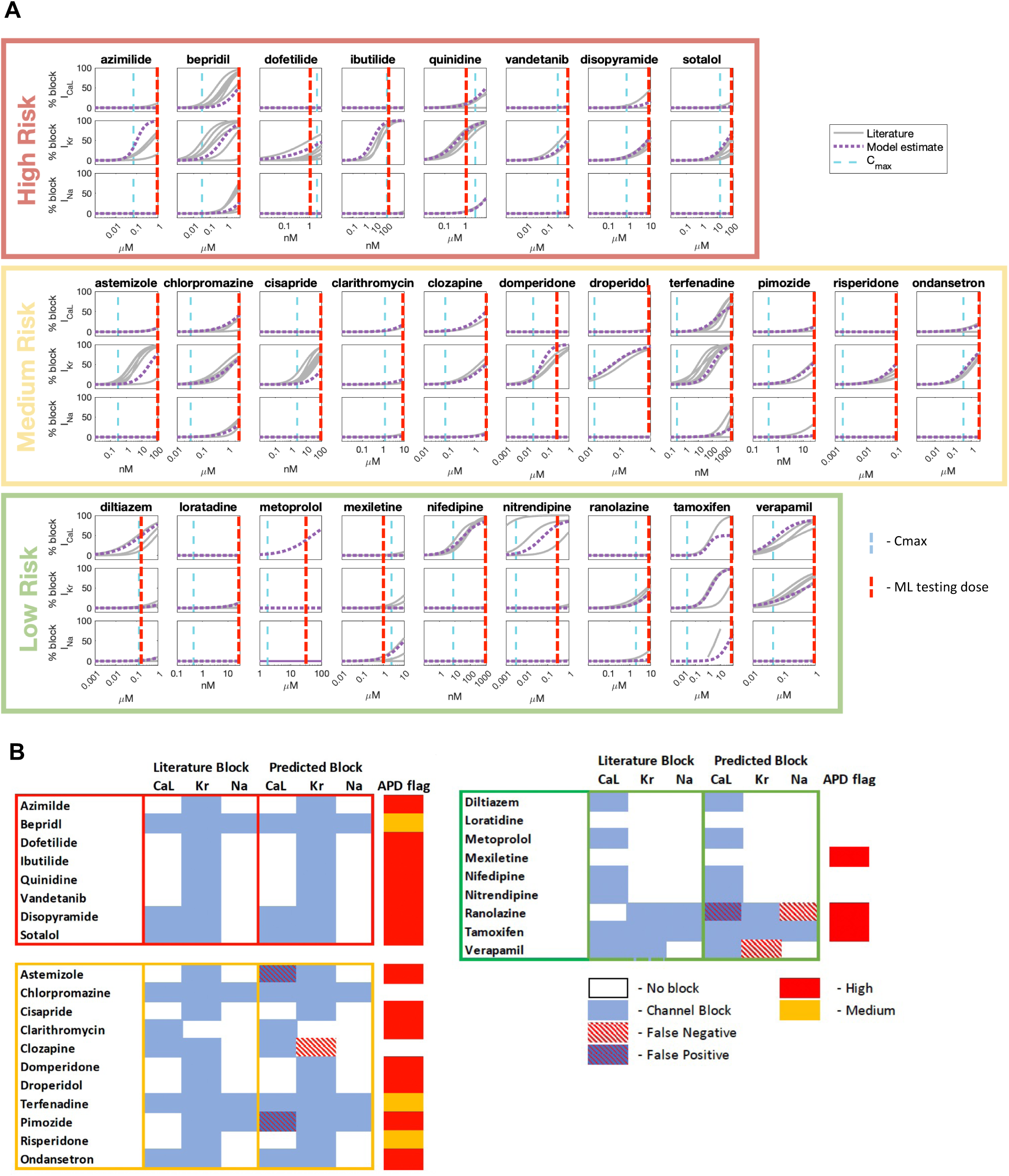
Modelling and Data Science results. **A**) Computational model determined block% of the L-type Calcium Channel (CaL), the hERG potassium channel (Kr), and the fast sodium channel (Na) for each compound (dotted purple line), which were compared to literature derived values (gray lines). The C_max_ for each compound is depicted with a vertical blue line, while the dose at which the machine learning algorithm was applied is depicted with a vertical red line. **B**) ML test results for each compound, used to determine if >10% block was present at the dose indicated in panel A. Expected block from literature is compared to block from the ML test results, with false positives and false negatives indicated. These are also compared to the AP analysis results in the final column.

### Machine learning channel block detection

Fitting of experimental data to biophysical models can be computationally difficult and time consuming. Therefore, we also applied a low computational cost machine learning algorithm as an alternative approach to mechanistically investigating channel block from action potential and motion biomarkers. The results are presented in **Figure 4B**, where a trained ML model was used on waveforms to predict if blocking of specific channels was greater than 10%. For each compound the computational model determined the % block of 3 channels: the L-type Calcium Channel (CaL), the hERG potassium channel (Kr), and the fast sodium channel (Na). Results (*predicted*) were compared to literature values (*literature block*) to look at accuracy and where false negatives and false positives occurred. Results indicate very good model accuracy against these heterogeneous measurements, with only 6 incorrect predictions out of 84 (3 ion channels for 28 compounds). Three failure modes were identified. The first is a false positive of calcium block, in the case of astemizole and pimozide, where the model predicts a block of >10%, but the literature values show blocks lower than 10% at the doses tested (red dashed lines). The second is for ranolazine, which predicted a calcium block instead of a sodium block, both of which could lead to the same phenotypic result of decreasing APD. The third is for compounds with significant block in both hERG and L-type Ca channels, verapamil and clozapine, where the calcium block was detected, but not the hERG block.

## Discussion

Over the last decade, hIPSC-CMs have become a standard in NAMs in assessing the arrhythmogenic potential of drug compounds,^15-18^ largely through the examination of potential or calcium dynamics^19, 20^ under the effect of drugs, although more extensive side effects have been studied using other biomarkers such as dynamic motion analysis.^21-23^ Most of these studies have used 2D plate formats to assess metrics,^7^ where hiPSC-CMs form a connected, but disorganized monolayer. However, 3D micromuscles have been shown to have improved predictive value over 2D plate formats^24^ with greater translatability due their improved maturity and coordinated tissue responses.^8, 9, 25, 26^

Drug induced APD prolongation is a main criterion for proarrhythmia risk flagging of new compounds. Particularly, the separation between C_max_ and doses leading to APD prolongation is critical to safety classification as it describes the relative risk for beating abnormalities in clinical applications. 10-fold C_max_ is often used as a safety margin cut-off in the development pipeline. A safety margin <10 will typically be considered as a “no-go” criterion and components will not move further in the development. Compounds with a >30-fold margin are typically considered safe.^27^

In addition to direct *in vitro* measurement of action potential, computational approaches have also played an important role in contextualizing measurements, and are also a part of the CIPA framework.^28, 29^ These biophysical models simulate the flow of ions across the cell membrane, and how altering this flow can change the action potential dynamic and lead to arrhythmia. Considerable work has gone into making improved and useful models of hiPSC-CMs^30^ to help understand and predict how changes measured in these cells can be interpreted into drug effects,^31, 32^ and how these changes can be expected to map to mature cardiomyocytes.^33, 34^ Finally, AI is currently taking a prominent role, helping to correlate how signals in hiPSC-CMs NAMs connect to clinical classifications.^35^

Here we connect these three approaches via the use of a high throughput cardiac microtissue model for arrhythmia risk prediction, where we layer contraction and electrophysiological data acquired using dose escalation studies with biophysical computational models and basic AI algorithms to illuminate drug risk profiles. We demonstrate the basic approach of evaluating APD changes is consistent with previous studies, capturing high risk compounds with known hERG block; however, this *in vitro* analysis alone can miss multiple aspects of channel block dynamics that may inform drug risk.

APD analysis alone was a reasonable metric that continues to give clarity on compound risk. By using a tiered risk assessment approach of changes in APD, we were able to identify clear risk signals for all high risk compounds close to the clinical C_max_ values. Either strong prolongation, the generation of arrhythmia, or both, was observed in this panel for 6/7 of the compounds, and the secondary risk profile flagged the last compound, bepridil. The *in silico* results further contextualize this, where both the cellular modeling and the AI pipeline show a more complex case for the compound with reduced potential risk, instead of simple hERG block near the C_max_. For other compounds, the combined modelling and AI approach gives greater contextualization of APD results. For intermediate compounds, we see risk signals in APD analysis for most of the compounds, and more complex blocking scenarios in the cases where no risk was flagged. For chlorpromazine, where multiple channels are blocked, there is no risk flagged by only APD. In addition, we observed the first instance of simultaneous block of both calcium and potassium in nearly equal levels, which can confound both APD analysis and the AI protocol as the hERG effect can be completely hidden by a concurrent change in calcium in these in vitro test systems.

One of the more interesting findings in this work is the extent that sodium block is a confounder to action potential based risk assessment. We see that the majority of the missed predictors from APD analysis alone have sodium block present, and this is apparent in the low risk compounds, where mexiletine, ranolazine, and tamoxifen all flag for APD risk. In each of these cases, we see a concurrent signal of increase in APD combined with cellular quiescence, and further study is needed to determine if this confounding result is specific to this cell type, as the same trends are seen in other studies using hiPSC-CM,^7^ or if this is a general trend.

Overall, this work establishes a high-throughput 3D cardiac micromuscle platform that integrates electrophysiological and mechanical readouts with computational biophysics and AI-based classification to improve proarrhythmia risk prediction. Using the CiPA compound panel as a reference point, we demonstrate that this layered *in vitro/in silico* approach robustly captures canonical action potential based risk signatures while also revealing complex MICE that are not detectable through APD analysis alone. In particular, the combined analyses illuminate cases in which sodium or multi-channel block confound traditional interpretation, and they highlight scenarios, such as simultaneous L-type calcium and hERG block, where integrated modeling materially improves mechanistic understanding.

By bridging advanced tissue engineering with quantitative modeling, this platform provides a more physiologically relevant and mechanistically transparent assay for compound cardiotoxicity assessment. The results support the potential of high-throughput analysis of 3D hiPSC-CM microtissues combined with computational models as a next-generation NAM for regulatory and preclinical workflows, improving the capacity to resolve nuanced drug effects that impact cardiac safety. Together, these findings demonstrate a practical, scalable, and information-rich framework that meaningfully advances predictive cardiotoxicology.

### Limitations of the study

The study was performed using a single, well characterized, healthy hiPSC-CM line. Previous studies on the CiPA drug panel have reported a consensus in results with minor line-dependent differences. To reflect a broader range of drug responses in a population, multiple and diverse cell lines should be used. Additionally, cell lines with known genetic mutations are needed to predict drug risks in subpopulations with specific genetic risks. Although a defined dose escalation protocol was employed in this study, our conclusions are limited to this acute exposure scenario. Any chronic, repeated or combined exposure effects need to be elucidated in additional experiments, where the cardiac NAM is suitable for these types of follow-up studies. Similarly, co-morbidities, which are known to increase arrhythmogenic drug risk, have been omitted in the present study, but could be modeled in follow-up experiments.

## Acknowledgements

We thank Dr. Mary West for support and access to the QB3 Cell & Tissue Analysis Facility/High-Throughput Screening Facility (UC Berkeley, California) and Dr. Gabriel Neimann for his support with confocal imaging. The work in this grant was funded by an NIH SBIR grant (R44TR004250) from the National Center for Advanced Translational Science (NCATS).

## Author contributions

Conceptualization, S.T.W., V.C., and K.E.H.; Methodology, V.C., S.T.W., A.R., B.K., H.K., K.H.J. and A.T.; Software, H.K., S.T.W., K.H.J. and A.T.; Formal Analysis, V.C., S.T.W., K.H.J. and A.T.; Investigation, V.C., A.R., K.H.J., B.K., H.K., B.S. and A.T.; Data Curation S.T.W., K.H.J. and A.T., Writing – Original Draft, V.C. and S.T.W.; Writing – Review & Editing, V.C., A.R., K.H.J., B.K., H.K., B.S., K.E.H., A.T. and S.T.W.; Visualization, V.C., S.T.W. and A.R.; Resources, S.T.W., A.T. and K.E.H.; Supervision, S.T.W.; Funding Acquisition, K.E.H. and S.T.W.

## Competing interests

K.E.H., H.K., K.H.J. and A.T. have a financial relationship with Organos Inc., and hence may benefit from the commercialization of the results of this research. In addition, K.E.H, S.T.W, and V.C have applied for patent protection for the described multiplexed high throughput test system.

## Declaration of generative AI and AI-assisted technologies

During the preparation of this work the authors used ChatGPT 5 [Large language model] in order to refine their original wording and grammar. After using this tool, the authors reviewed and edited the content as needed and take full responsibility for the content of the published article.

## Supplemental information

Table S1. Excel file containing drug compound details, related to **Table 1**

Supplementary Figure 1.

STAR* **Methods**

## STAR*Methods

**STAR\*Methods**
| **REAGENT or RESOURCE** | **SOURCE** | **IDENTIFIER** |
| --- | --- | --- |
| ***Chemicals, Peptides, and Recombinant Proteins*** |  |  |
| plating media | Fujifilm Cellular Dynamics | M1001 |
| maintenance media | Fujifilm Cellular Dynamics | M1003 |
| PBS | Corning | 21-031-CV |
| penicillin-streptomycin | Sigma-Aldrich | P4333 |
| Tamoxifen | Sigma-Aldrich | T2859 |
| Nitrendipine | Sigma-Aldrich | N144 |
| Pimozide | Sigma-Aldrich | P1793 |
| Loratidine | Sigma-Aldrich | L9664 |
| Droperidol | Sigma-Aldrich | D1414 |
| Chlorpromazine hydrochloride | Sigma-Aldrich | C8138 |
| Clozapine | Sigma-Aldrich | C6305 |
| Risperidone | Sigma-Aldrich | R3030 |
| Ranolazine | Sigma-Aldrich | PHR2676 |
| Clarithromycin | Sigma-Aldrich | A3487 |
| Sotalol hydrochloride | Sigma-Aldrich | 1617408 |
| Ondansetron Hydrochloride | Sigma-Aldrich | 1478582 |
| Domperidone | Sigma-Aldrich | D122 |
| Diltiazem, Hydrochloride | Sigma-Aldrich | 309866 |
| Mexiletine hydrochloride | Sigma-Aldrich | M2727 |
| Vandetanib | Sigma-Aldrich | SML2983 |
| Astemizole | Sigma-Aldrich | A2861 |
| Ibutilide hemifumarate salt | Sigma-Aldrich | I9910 |
| Azimilide dihydrochloride | Sigma-Aldrich | SML0353 |
| Bepridil hydrochloride | Sigma-Aldrich | B5016 |
| dofetilide | Sigma-Aldrich | PZ0016 |
| Quinidine free base anhydrous | Sigma-Aldrich | Q3625 |
| Cisapride monohydrate | Sigma-Aldrich | C4740-50mg |
| terfenadine | Sigma-Aldrich | T9652 |
| (±)-Metoprolol (+)-tartrate salt | Sigma-Aldrich | M5391-1G |
| nifedipine | Sigma-Aldrich | PHR1290-1G |
| (±)-Verapamil hydrochloride | Sigma-Aldrich | V4629-1G |
| disopyramide | Sigma-Aldrich | D2920000 |
| dimethyl sulfoxide (DMSO) | Sigma-Aldrich | D8418 |
| voltage sensitive dye <sup>a</sup> | Photoswitch Biosciences | PhoS1 |
| 4% paraformaldehyde solution | Thermo Fisher | AAJ61899AK |
| fetal bovine serum | Sigma Aldrich | 1943609-65-1 |
| Triton X-100 | Sigma Aldrich | X100 |
| Alexa Fluor 488 phalloidin | Sigma Aldrich | A12379 |
| anti-cardiac isoform troponin T | Epredia | MS-295-P |
| anti-mouse Alexa Fluor 594 | Thermo Fisher | A11005 |
| DAPI | Sigma Aldrich | MBD0015 |
| **Specialty labware** |  |  |
| 384 well open bottom plates | GraceBioLabs | 206384 |
| adhesive film | Research Products International | 202503 |
| TOPAS thermoplastic elastomer (cyclic olefin copolymer) | STRATEC Consumables GmbH, Austria | TPE E140 (custom form) |
| glass coverslip | Ibidi GmbH, Germany | 10812 |
| **Experimental models: Cell Lines** |  |  |
| Human iPSC line (healthy donor) | Fujifilm Cellular Dynamics | Cat# R1007 |
| **Software and algorithms** |  |  |
| Harmony | Perkin Elmer | ver. 4.9.2 |
| Imaris Bitplane | Oxford Instruments | 9.2 |
| ImageJ2 | NIH | ver. 2.14 |
| Prism 10 for macOS | GraphPad Software, LLC | ver. 10.4.1 (532) |
| MATLAB | Mathworks | R2024B |
| MPS-Motion | https://github.com/ComputationalPhysiology/automatic-motion-estimation/ | V1.0.0 |

**a)** identical to BeRST-1 dye<sup>38</sup>

### Plate Fabrication

Plates for 3D microtissue formation were fabricated from ANSI compatible 384 well plates with open bottoms (206384, GraceBioLabs). The plates come with an adhesive film that was used to our mount custom substrates to the plate bottom. Substrates were made from TOPAS thermoplastic elastomer (TPE) E-140, which is a flexible semicrystaline cyclic olefin copolymer (COC) with excellent optical clarity and low water permeability, but high oxygen permeability. The TPE was injection molded thermoplastic by STRATEC Consumables GmbH, Austria, based on our CAD designs (**Fig. 1D**). The injection molded TPE was delivered as disks (11.9cm diameter, 1.3 mm thickness) with 2 strips (2.5x7.5cm) (**Fig. 1B**), each containing microfeatures for 5x16 wells. TPE strips were cut from the disks, rinsed with 70% ethanol, blow dried and disinfected under UV light for 30min before assembly inside a biosafety cabinet. Four TPE strips were manually aligned and adhered to each 384 well plate yielding a total of 320 wells per plate (**Fig. 1A**). Each well comprised 4 micromuscle troughs (1400x200x150µm length x width x depth, that allow for self-assembly of hiPSC-derived cardiomyocytes (hiPSC-CM) into uniaxially contracting muscle fibers (**Fig. 1E**). The finished plate was again subjected to UV disinfection. A sterile adhesive film (202503, RPI Research Products International) was applied to seal the top of the plate and maintain sterility until use.

### Cell Loading

The plate seal film was cut and removed from the wells to be used for cell loading. A sterile plate lid (3099, Corning) was added. Cardiomyocytes (hiPSC-CM) and media were purchased from Fujifilm Cellular Dynamics (Donor#: 01434, Cat# R1007). The cells were thawed from liquid nitrogen cryo-storage according to the vendor protocol. The cell pellet was suspended in plating media (M1001, Fujifilm Cellular Dynamics) supplemented with penicillin-streptomycin (P/S) (P4333, Sigma) to 35,000 viable cells per 2.5µL (14x10^6^ cells/mL). To remove air bubbles from the microfeatures, each well was briefly rinsed with 70% ethanol, followed by a single wash step with sterile PBS (21-031-CV, Corning). PBS was aspirated and 2.5µL of singularized cell suspension were carefully placed on top of the microfeatures in the center of each well (**SuppFig. 1A,B**). The droplet was not allowed to touch the well wall on any side to keep the cells concentrated at the microfeatures. The cell suspension was gently mixed after every 4 ^th^ well to ensure homogeneous cell distribution. Once cell droplets were dispended to each well, the cells were allowed to sediment for 3 minutes. A centrifugation step (1min at 100xG) was applied to pack the cells into the micromuscle troughs while washing residual cells off the plateau area between the micro-troughs. Excess cell suspension was aspirated from the corner of each well. Using a multichannel pipette, 50µL plating media were added to each well and another centrifugation step (1min at 100xG) ensured that the media reached the bottom of each well. After 48h the media was changed to maintenance media (M1003; Fujifilm Cellular Dynamics) with 1% P/S and replaced every 2 to 3 days. The samples were maintained for 6-15 days to allow for tissue formation (**Fig. 1E-H, SuppFig. 1C**) before drug testing. Some of the thinner tissue strands eventually broke under the continuous mechanical load. Up to four independently beating cardiac micromuscles formed in each well. On the day before a drug test, samples were stained with 500nM voltage sensitive dye (PhoS1; Photoswitch Biosciences) in maintenance media overnight. Media needed for drug preparation was placed in the cell culture incubator over night to equilibrate temperature and pH. On the day of drug testing staining media was replaced with 50µL assay media (M1003 + 1% P/S + 50nM PhoS1).

### Drug preparation

All compounds were purchased in powder format and dissolved in dimethyl sulfoxide (DMSO) (D8418, Sigma-Aldrich) to create stocks for screening. Supplementary Table 1 details the drug compounds and stock preparation. Each drug was tested in a 5-step dose-escalation study (Table 1) where the next higher dose was consecutively added to the same well. DMSO served as a solvent control and was added up to 0.1% in a log-scale dilution. All drug concentrations were freshly prepared in assay media so that the sequential addition of 5µL per well yielded the desired final drug dose.

**Table 1:**
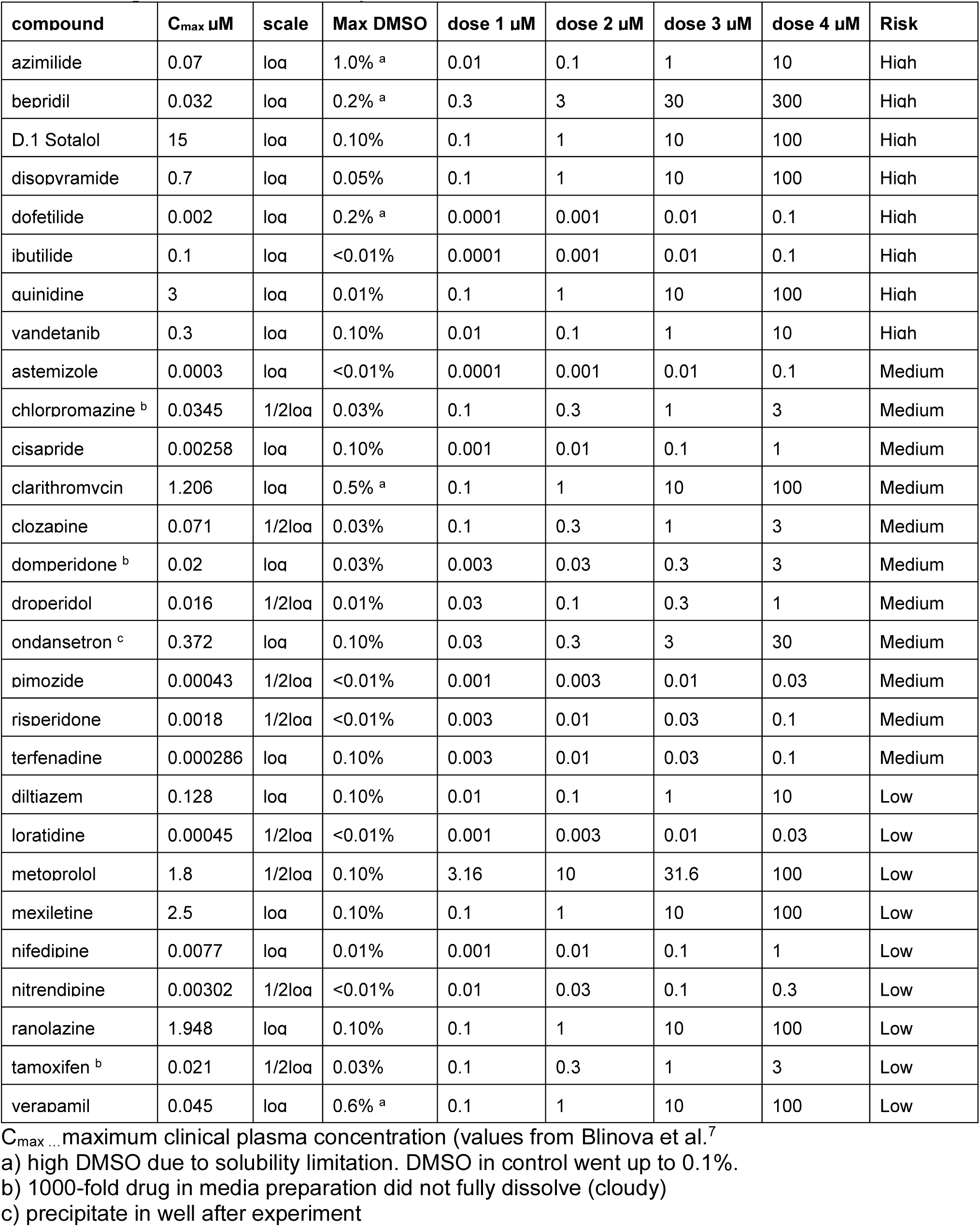
Drug doses used for compounds tested.

### Drug application and image acquisition

All imaging was performed inside the incubation chamber of an ImageXpress Micro (Molecular Devices) microscope set to 37°C, 5% CO_2_, humidified. A 60min equilibration time was allowed before baseline traces were recorded. Every compound was tested in 3 to 5 independent wells containing up to 4 micromuscles each. Each well started out with 50µL assay media to which 5µL of increasing drug concentration were added at each step of the dose-escalation study. Imaging was performed at baseline and after application of each drug dose. Every dose was incubated for 20min before acquisition was started. Image acquisition took 34s per well, and after acquisition of all wells at a given dose, the next higher dose was applied to all wells. A total of up to 80 wells were analyzed per experiment resulting in incubation times of 20 to 65 min for the first vs last well to be imaged. To reach an average incubation time of about 42min for each compound the replicate wells were distributed over the plate. Brightfield (14% light power) and Cy-5 fluorescence (1.5% laser power) were recorded in each well for 8s at 50fps via a 4x objective with 2x2 binning.

### Immunofluorescence

Tissues were washed twice with PBS, then fixed in 50µL per well 4% paraformaldehyde solution in PBS (AAJ61899AK, Thermo Fisher) for 20 min at room temperature and washed three times with blocking buffer consisting of PBS with 5% fetal bovine serum (1943609-65-1, Sigma Aldrich) and 0.1% Triton X-100 (X100-100ML, Sigma Aldrich). Staining solution was prepared in blocking buffer with 1:400 Alexa Fluor 488 phalloidin (A12379, Thermo Fisher) 66 μM stock in DMSO, 1:100 anti-cardiac isoform troponin T antibody (MS-295-P, Epredia) and 1:1000 DAPI (MBD0015-1mL, Sigma Aldrich). Samples were incubated with 50µL staining solution for 2h on a rocker at room temperature followed by overnight incubation at 4°C. After three washes in blocking buffer the secondary antibody anti-mouse Alexa Fluor 594 (A11005, Thermo Fisher) was added at 1:200 (same incubation times and temperatures as primary antibodies) followed by three washes in blocking buffer. For imaging the TPE substrates were removed from the 384 well GraceBioLabs plates and mounted upside-down on a 170um thick glass coverslip (10812, ibidi) in blocking buffer. Image acquisition was performed using an Opera Phenix (Perkin Elmer) confocal microscope using 40x magnification and 1.15 NA objectives, with DAPI(375/456nm), FITC (488/522nm), and TxRed (562/599nm) filters. Video analysis and processing was performed using Harmony (Perkin Elmer, ver. 4.9.2), Imaris Bitplane (Oxford Instruments, ver. 9.2) and ImageJ2 (NIH, ver. 2.14)

### Data processing

All acquired videos were automatically processed to obtain a range of relevant electrical and motion biomarkers (**Fig. 2A**). First, a video was automatically segmented into individual regions of interest encompassing a single micromuscle. These ROIs were there processed using automated scripts to obtain beat averaged motion (displacement) and voltage traces for individual tissues.^39^ QC criteria were applied on the initial (no dose) fluorescence and motion traces, eliminating individual troughs with insufficient tissue present for analysis. From these average traces, a range of biomarkers were calculated, including APD80 and beat rate, and also measures of the motion such as displacement duration at 50% and 80% (DD50 and DD80 respectively) for each tissue across the dose escalation. Beat rate changes inherently cause a change in APD. To distinguish beat rate effects from actual action potential modulation, beat rate corrected data (cAPD) was applied. Different algorithms have been published^40^ and here we use the Fridericia formula cAPD = APD * BR^1/3^.^41^ These automatically computed biomarkers were then manually inspected together with the averaged traces, and if biomarker computation had been confounded by arrhythmias, the respective values were excluded from further analysis, and the arrhythmias noted. In this stage, EADs were also manually identified by analysis of the voltage traces.

### Computational model of cellular biophysics

We used an ODE based computational biophysical model of hiPSC-CMs based on previous work^31^ adjusted to represent spontaneously contracting cells. Here, a dynamic intracellular Na^+^ concentration is incorporated,^42^ and the mechanical contraction of the cells was simulated by incorporating a cross bridge model,^37^ with mechanical parameters unchanged except that *k*_on_ is increased from 0.05 *µ*M^−1^ms^−1^ to 0.45 *µ*M^−1^ms^−1^ and *Pcon*_titin_ is increased from 0.002 to 0.018. All modified parameter values of the base model are as reported in Table 2. The resulting ODE system is solved numerically using the MATLAB ode15s solver.

**Table 2:**
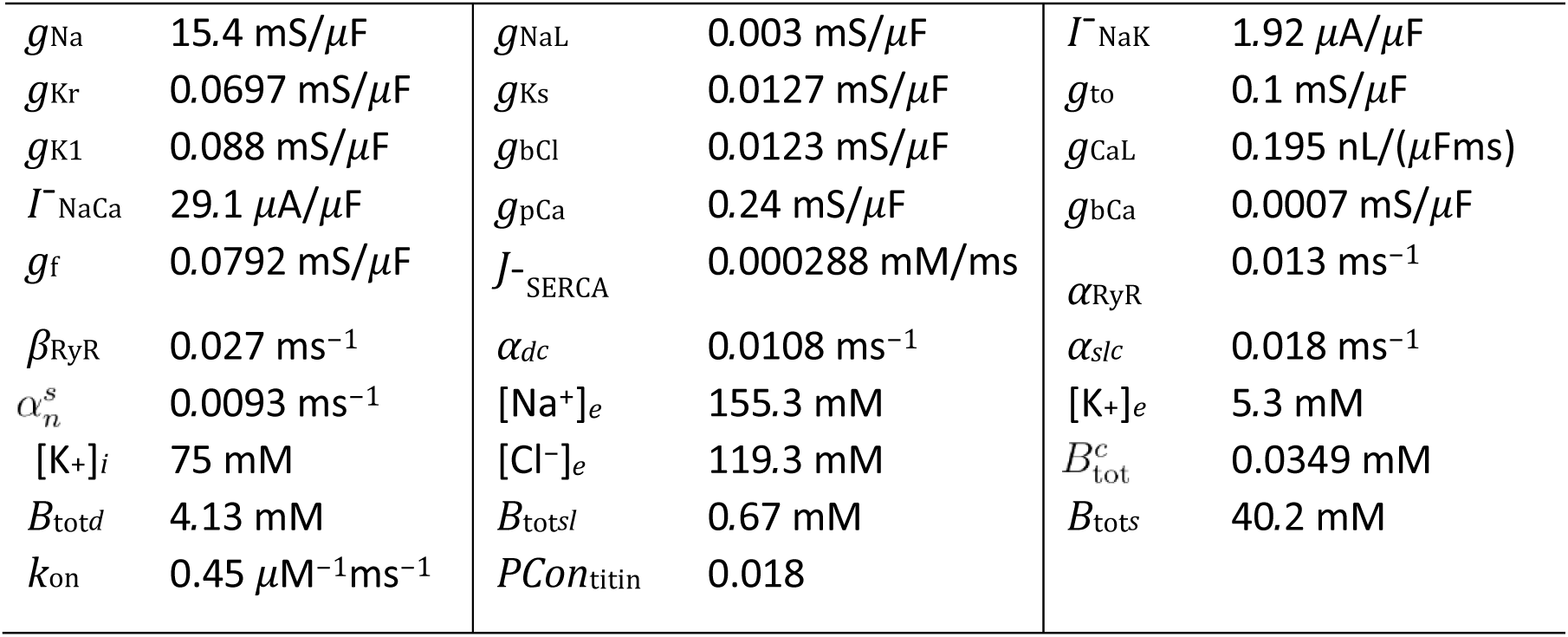
hIPSC-CM Model Parameterization. Parameter values used in the computational model. The remaining parameter values are as previously published.^36, 37^

### Representation of drug effects

The dose dependent effect of a drug compound on a model parameter, *p*, is represented as previously,^43^ i.e., by

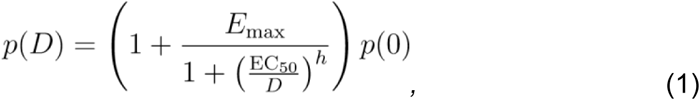

where *p*(*D*) is the parameter value in the presence of the drug dose *D*, *p*(0) is the default parameter value, *h* is a Hill coefficient, EC_50_ is the half maximal effective concentration, and *E*_max_ is the maximum effect of the drug in terms of percentage block. For the compounds in this study, we consider drug effects on the membrane model parameters *g*_CaL_, *g*_Kr_, *g*_Na_, *g*_NaL_, *g*_K1_, *g*_bCa_, *g*_f_, and *Ī*_NaCa_ representing adjustment of the L-type calcium channel current, the hERG rapidly activating potassium current, the fast sodium current, the late sodium current, the inward rectifying potassium current, the background calcium current, the hyperpolarization activated funny current, and the Na^+^/Ca^2+^ exchanger current, respectively.

### Estimation of dose-dependent drug effects

For each drug and model parameter, the drug parameters *h*, *E*_max_ and EC_50_ were estimated based on values from an extensive literature survey.^44-100^ and manual tuning performed such that biomarker changes estimated by the model align well with the relative biomarker changes observed in the cardiac micromuscle. In cases where the literature references did not provide values for *h* or *E*_max_, these drug parameters are assumed to be 1 and −1, respectively. The biomarkers considered in the comparison between model and measurements were beat rate, the maximum displacement amplitude (Peak disp.), and the duration at 50% and 80% of the action potential (APD50 and APD80) and the displacement curves (DD50 and DD80). All biomarkers are considered as relative changes to the baseline (no-drug) case. In the model, the beat rate, APD50 and APD80 are computed from the change in membrane potential whereas the Peak disp., DD50 and DD80 are computed from the change in sarcomere length. After a parameter change is applied, the simulations are run for 240 seconds to allow reaching dynamic equilibrium before measuring the updated biomarker values.

### Automatic identification of *I*_CaL_, *I*_Kr_, and *I*_Na_ block

To automatically estimate whether a given compound that blocked the *I*_CaL_, *I*_Kr_, or *I*_Na_ currents, a random forest based classification algorithm was applied. For a given drug dose, we computed the biomarker changes compared to the baseline case for four biomarkers: APD80; beat rate; Peak disp; and the upstroke time (*t*_up_). We defined *t*_up_ as the time from the membrane potential is halfway between the minimum and maximum values during the upstroke until it reaches the value 20% below the maximum value. The changes in these biomarkers in response to the drug was subsequently classified as ‘increase’, ‘unchanged’, or ‘decrease’. The thresholds between when a biomarker was ‘unchanged’ or not was set to 5% for APD80, 20% for the beat rate, 50% for *t*_up_, and 7.5% for Peak disp.

The classified changes in these four biomarkers were then used as input in a random forest classifier. The output of the classifier was an estimate of which, if any, of the three currents, *I*_CaL_, *I*_Kr_, or *I*_Na_, were blocked by the drug. The random forest classifier was constructed using the TreeBagger function in MATLAB with 100 trees. The training data for the classifier was based on 70,000 simulations of the computational hiPSC-CMs model for different randomly drawn combinations of block of the three considered currents. Blocking of currents of less than 10% were classified as no block. To better represent the experimental data, random noise was added to the biomarkers computed from the simulation results. The noise was drawn from a normal distribution with expectation equal to zero and standard deviation equal to half of the threshold values used to determine whether a biomarker was unchanged or not (see, above).

We applied this automatic identification algorithm to the highest considered drug dose of each drug (excluding doses so high that the cells stopped beating or displayed early after depolarizations) for a single selected tissue exposed to each drug. The considered tissues were selected manually such that the displayed biomarker changes of the tissue aligned as well as possible with the biomarker changes observed in computational model simulations of the drug, using the effects found in literature from the above survey.

## Notes

https://github.com/karolihj/drug-identification-2026

