## supplementary information for "Drug Proarrhythmic Evaluation in a High Throughput Cardiac New Approach Methodology"

Contents:

- Supplementary Figure 1
- Supplementary Table 1

### Supplementary Figure 1

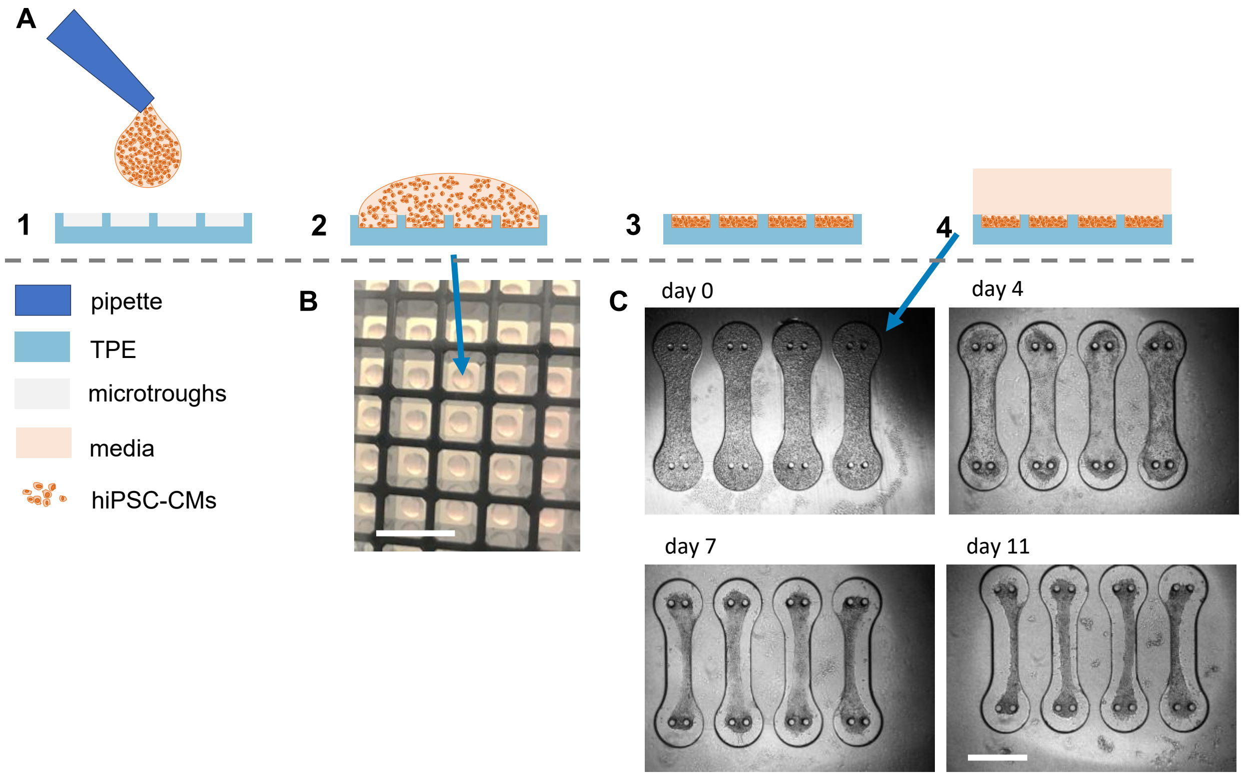


Supplementary Figure 1: Cell loading into the microtroughs: A) 1) A 1.5µL droplet of cell suspension is carefully placed on top of the microstructures in the center of each well. 2) The cell suspension covers the micropatterned area. 3) After cell sedimentation and centrifugation cells are packed into the microtroughs while the plateau region is cleared of excess cells and media. 4) Fresh media is carefully added to the well. B) Photo of a section of a 384 well plate with loading droplets. Scale bar represents 5mm. C) Microtissue formation in microtroughs after loading with cell suspension. Scale bar represents 500µm.

### Supplementary table 1: Drug compound overview

| Drug | Risk group [a] | Compound | Product number [b] | MW (g/mol) | CAS number | Stock (mM) [c] |
|---|---|---|---|---|---|---|
| azimilide | H | Azimilide dihydrochloride | SML0353 | 530.88 | 149888-94-8 | 1 |
| bepridil | H | Bepridil hydrochloride | B5016 | 403 | 68099-86-5 | 10 |
| D,1 Sotalol | H | Sotalol hydrochloride | 1617408 | 308.82 | 959-24-0 | 100 |
| disopyramide | H | disopyramide | D2920000 | 339.47 | 9/5/37 | 200 |
| dofetilide | H | dofetilide | PZ0016 | 441.56 | 115256-11-6 | 10 |
| ibutilide | H | Ibutilide hemifumarate salt | I9910 | 442.61 | 122647-32-9 | 10 |
| quinidine | H | Quinidine free base anhydrous | Q3625 | 324.42 | 56-54-2 | 30 |
| vandetanib | H | Vandetanib | SML2983 | 475.35 | 443913-73-3 | 10 |
| astemizole | M | Astemizole | A2861 | 458.57 | 68844-77-9 | 10 |
| chlorpromazine | M | Chlorpromazine hydrochloride | C8138 | 355.33 | 69-09-0 | 10 |
| cisapride | M | Cisapride monohydrate | C4740-50mg | 483.96 | 260779-88-2 | 10 |
| clarithromycin | M | Clarithromycin | A3487 | 747.95 | 81103-11-9 | 20 |
| clozapine | M | Clozapine | C6305 | 326.82 | 5786-21-0 | 10 |
| domperidone | M | Domperidone | D122 | 425.91 | 57808-66-9 | 10 |
| droperidol | M | Droperidol | D1414 | 379.43 | 548-73-2 | 10 |
| ondansetron | M | Ondansetron Hydrochloride | 1478582 | 365.85 | 103639-04-9 | 30 |
| pimozide | M | Pimozide | P1793 | 461.55 | 2062-78-4 | 10 |
| risperidone | M | Risperidone | R3030 | 410.48 | 106266-06-2 | 10 |
| terfenadine | M | terfenadine | T9652 | 471.67 | 50679-08-8 | 10 |
| diltiazem | L | Diltiazem, Hydrochloride | 309866 | 451 | 33286-22-5 | 10 |
| loratidine | L | Loratidine | L9664 | 382.88 | 79794-75-5 | 30 |
| metoprolol | L | (±)-Metoprolol (+)-tartrate salt | M5391-1G | 684.82 | 56392-17-7 | 100 |
| mexiletine | L | Mexiletine hydrochloride | M2727 | 215.72 | 1/4/70 | 100 |
| nifedipine | L | nifedipine | PHR1290-1G | 346.33 | 21829-25-4 | 10 |
| nitrendipine | L | Nitrendipine | N144 | 360.36 | 39562-70-4 | 10 |
| ranolazine | L | Ranolazine | PHR2676 | 427.54 | 95635-55-5 | 100 |
| tamoxifen | L | Tamoxifen | T2859 | 371.51 | 10540-29-1 | 10 |
| verapamil | L | (±)-Verapamil hydrochloride | V4629-1G | 491.06 | 0152-11-4 | 10 |

a) risk groups: L … low risk, M … intermediate risk, H … high risk
b) all compounds purchased from Sigma-Aldrich
c) all stocks prepared in DMSO and stored at -80°C
